# Short-Term Combined Tat-Beclin1 and Endurance Training Improves Age-Related Decline in Physical Function in Male Mice

**DOI:** 10.64898/2026.05.07.723527

**Authors:** Thuan Thien Tchen, Shakibur Rahman, Thaysa Ghiarone, Lynn A. Spruce, Hossein Fazelina, Elizabeth M. Brown, Charalampos Papachristou, Sue C. Bodine, Vitor A. Lira, Kleiton A. S. Silva

**Author notes:** Corresponding Author: Kleiton A. S. Silva, PhD, Cooper Medical School of Rowan University 401 S Broadway, Camden, NJ, USA 08103.

## Abstract

Autophagy is a hallmark of aging, but autophagy-related proteins have not been exclusively targeted to attenuate the progressive decline in physical function associated with aging. Here, we combined Tat-Beclin1, an autophagy agonist, and endurance training to determine whether Tat-Beclin1 enhances exercise adaptation in old male mice. Tat-Beclin1 was administered intraperitoneally (TB group, 15 mg/kg, 2x/week) as a standalone therapy, or in combination with endurance training (TB+Exe group, 70% of maximal running speed 3x/week) for 1 month in 23-month-old male C57BL/6J mice. Control groups were age-matched cage controls and exercise-only groups. Animals were assessed for grip strength, endurance capacity on a treadmill, and balance and coordination on a rotarod. Gastrocnemius/plantaris (G/P) and tibialis anterior muscles were harvested for western blotting, myofiber typing, and proteomic profiling (G/P only). TB+Exe led to significant increases in grip strength, endurance capacity, and balance and coordination performance beyond those observed in the TB and Exe groups alone. Autophagy markers, including Beclin1, the LC3B-II/I ratio, and p62, did not differ among groups. A proteomic analysis of the G/P muscle revealed that TB upregulated biological processes involved in muscle contraction and adaptation, whereas TB+Exe increased mitochondrial bioenergetic processes and, surprisingly, upregulated acute inflammatory responses, including proteins such as haptoglobin and orosomucoid-1. We conclude that combining Tat-Beclin1 and endurance training may represent a new approach to attenuate aging-related decline in physical function.

**New & Noteworthy:** We show evidence that combining Tat-Beclin1 and endurance training (TB+Exe) resulted in greater improvements in physical function in 24-month-old male mice than either standalone therapy. We also show that TB+Exe upregulates traditional exercise-like biological processes and unexpectedly upregulates acute-inflammatory proteins (e.g., orosomucoid-1), which are thought to improve physical function in preclinical studies. Our study suggests that TB may be a new drug enhancing physical function, especially when combined with endurance training in old male mice.

## Introduction

Aging leads to a progressive decline in physical function, negatively impacting the quality of life of affected individuals (1–5). Given that the world population is aging at an alarming pace (6, 7), developing therapies improving overall physical function is essential for extending healthspan (8). Regular exercise is an effective approach for slowing aging-related deterioration in physical function, but limitations, including physical debilitation and age-associated diseases, prevent many older adults from adhering to an exercise program (9). Intriguingly, even among long-term exercise practitioners, the negative effects of aging may limit the benefits of exercise (1, 9). Several factors contribute to the development and progression of functional decline, including insulin resistance, mitochondrial dysfunction, chronic low-grade inflammation, impaired proteostasis and macroautophagy (hereafter referred to as autophagy), and cardiovascular dysfunction (10–12). In an attempt to identify a drug that attenuates aging-induced physical dysfunction, many pharmacological agents, including metformin, rapamycin, vitamin D, testosterone, SS-31, and MitoQ (3, 8, 13–15), have shown promising results in reducing aging-related physical dysfunction. Conversely, when testing some of these agents in combination with regular exercise, available data suggest that exercise adaptations are limited or blunted by the combinatory approach (16–19). Therefore, it is imperative to expand our understanding of the mechanisms that modulate aging-related physical dysfunction and to identify effective therapies that may enhance the benefits of exercise.

Autophagy is the primary cellular mechanism for degrading long-lived proteins and organelles and is essential for opposing physical dysfunction (20, 21). It has been postulated that autophagic activity declines with aging (12), making autophagy a suitable target to counteract age-related declines in physical function. Although agents that attenuate physical dysfunction may modulate autophagy, they do so by targeting molecules that are not constitutively embedded in the autophagy machinery (i.e., autophagy-related proteins). Beclin1 is a fundamental autophagy-related protein that coordinates autophagosome formation and fusion with lysosomes through its association with the class III phosphatidylinositol 3-kinase (PtdIns3K) complex (22). The role of Beclin1 in maintaining physical function has been demonstrated in a young Beclin1 haploinsufficient mouse model, in which Beclin1 deficiency blunted exercise-induced increases in capillary density and endurance capacity (23).

Additionally, in a transgenic mouse model with a BECN1^F121A^ knock-in mutation that activates autophagy by disrupting the Beclin1/BCL2 interaction, myofiber cross-sectional area was preserved in 20-month-old mice (24). These data demonstrate that genetically targeting Beclin1 is sufficient to alter exercise adaptation and improve aging-related muscle dysfunction.

Nevertheless, only one study has pharmacologically targeted Beclin1 (i.e., Tat-Beclin1) in a mouse model with sustained mTOR activation, showing that activation of autophagy improves mitochondrial function and prevents neuromuscular junction dysfunction (25). Tat-Beclin1 is a small permeable peptide derived from the Beclin1 protein (26). Mechanistically, Tat-Beclin1 binds to the Golgi-associated plant pathogenesis-related protein 1 (GAPR-1), a negative regulator of autophagy, releasing Beclin1 back into the PtdIns3K complex and triggering autophagy (26, 27). Yet whether Tat-Beclin1 serves as a pharmacological agent counteracting age-related physical dysfunction remains elusive.

In this study, we hypothesized that combining Tat-Beclin1 and endurance training in an alternating-day regimen would yield superior benefits in physical function assessments (i.e., grip strength, endurance capacity, and balance and coordination compared with single-mode treatments and would identify key molecules associated with improvements in physical function. We find that Tat-Beclin1 alone improves grip strength and endurance capacity, whereas endurance training increases only endurance capacity. Remarkably, combining these approaches further enhances grip strength, endurance capacity, and balance and coordination. Additionally, our proteomics analysis showed that, while Tat-Beclin1 upregulated several biological processes related to skeletal muscle contraction and adaptation, the combinatorial strategy increased mitochondrial bioenergetic processes and unexpectedly upregulated proteins involved in the acute inflammatory response, revealing a complex and novel regulatory mechanism that attenuates skeletal muscle aging. Taken together, our findings demonstrate that Tat-Beclin1 is a promising candidate for attenuating aging-related physical dysfunction, particularly when combined with endurance training.

## Materials and Methods

### Animals

Male C57BL/6J mice were purchased from the Jackson Laboratory (Bar Harbor, ME) at 22 months of age. After 2 weeks, at 22.5 months of age, animals were used for experimental purposes (i.e., physical function assessments). Upon arrival at the Cooper Medical School of Rowan University vivarium, animals were randomly housed in a group of 3-5 mice per cage.

Housing conditions included an enriched cage environment, a standard chow diet (5LG4 - LabDiet® JL Rat and Mouse /Irr 6F, Richmond, IN), and water ad libitum, with room temperature controlled at 21-22 °C, and light and dark cycles of 12 hours each. Mice were randomly divided into Control (N=9), Tat-Beclin1 (TB, N=9), exercise (Exe, N=9), and TB+Exe, N=8 (Figure 1).

**Figure 1.**
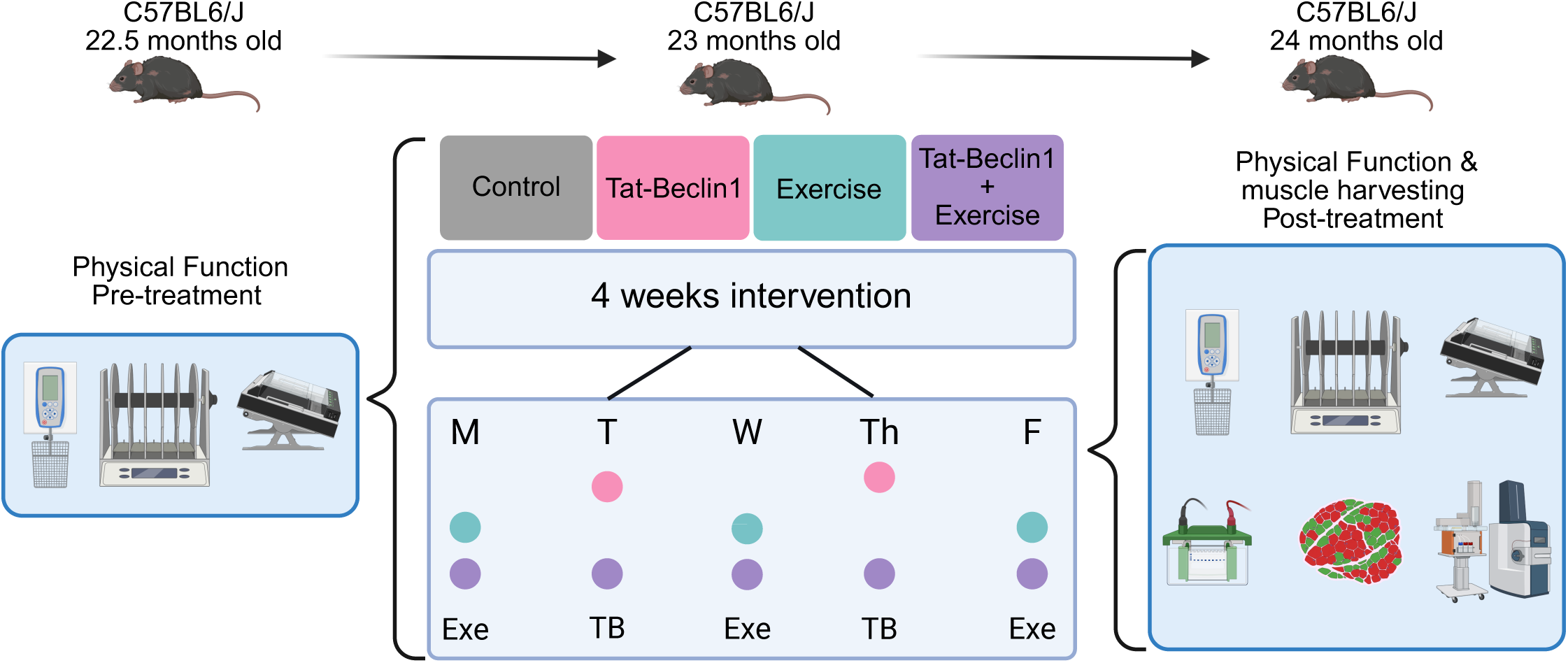
Experimental Design. Mice underwent physical function testing before the intervention, including a familiarization period followed by baseline testing. Twenty-three-month-old male C57BL6/J mice were assigned to Control, Tat-Beclin1 (TB), Exercise (Exe), and TB+Exe groups and completed a 4-week intervention in an alternating-day regimen. Exercise was performed three times per week (Monday, Wednesday, and Friday), and Tat-Beclin1 was administered two times per week (Tuesday and Thursday). After the intervention, mice underwent post-treatment physical function testing, and muscles were harvested for morphological, protein-expression, and proteomic analyses.

Forty-eight hours after the last Tat-Beclin1 injection (TB group) or the last exercise session (Exe and TB+Exe groups), all animals were euthanized between 1:00 PM and 5:00 PM by CO_2_ overdose followed by cervical dislocation. All animal protocols were performed under the Principles for the Utilization and Care of Vertebrate Animals Used in Testing, Research, and Training and approved by the Rowan University Animal Care and Use Committee (Protocol No. 2022-1316).

### Physical function

Two weeks after arrival, we conducted physical function measurements at the pre- and post-treatment periods. Animals were familiarized with a grip strength meter (Cat. No. 76-1068), a motorized treadmill (Cat. No. LE8710MTSAPC), and a rotarod (Cat. No. LE8205), all purchased from DSI/Panlab/Harvard Apparatus. For the grip strength experiment, mice were positioned with all four paws on the grid, and a designated investigator lightly but securely held each mouse by the base of the tail. After confirming that the animal had grasped the grid, the investigator gently pulled the animal back. The grip strength of the mouse’s four paws was recorded in Newtons (N). We conducted 5 trials per animal, allowing 1 minute of rest between trials (28). For the rotarod test, mice were habituated to the rotarod 24 hours before testing.

Briefly, animals performed three trials, each consisting of an accelerating rotarod set at a gradual speed (i.e., 4-40 rpm) for 5 minutes (29). The latency to fall (in seconds) was recorded. After completing the grip strength and rotarod assessments, animals were familiarized with a motorized treadmill as follows: day 1: stationary treadmill belt at 10% slope for 10 minutes; day 2: 5 m/min for 5 minutes at 10% slope; day 3: 5 m/min for 5 minutes at 15% slope; day 4: 5 m/min for 5 minutes at 20% slope; day 5: 5 m/min for 5 minutes at 25% slope; day 6: resting; and day 7: incremental test, which served as an evaluation of endurance capacity and for training prescription. The incremental test consisted of a warm-up period at 5 m/min for 3 minutes at a 25% slope; the speed was then increased by 5 m/min every 1 minute until exhaustion (30). Failure to maintain treadmill speed was characterized by contact with the tail or hindlimbs on the back of the acrylic lane, or by standing or sitting on the grid, which elicited a grid shock (0.2 V) for approximately 5 seconds. Endurance capacity was reported as time to exhaustion in seconds (3).

### Endurance training

At 23 months of age (Figure 1), animals in the Exe and TB+Exe groups underwent one month of endurance training. Although animals from the Control and TB groups did not participate in exercises, they were brought to the exercise room and positioned near the treadmill to account for stress associated with treadmill operation. Exercise sessions were held from 1:00 PM to 5:00 PM on Mondays, Wednesdays, and Fridays, for a total of 12 sessions.

Each session included a warm-up period of 5 m/min for 5 minutes, a training period of 33 minutes (during the first and second weeks) and 43 minutes (during the third and fourth weeks) at 70% of the maximum running speed, and a cool-down period of 5 m/min for 2 minutes. The treadmill slope was maintained at 5% in the first week and increased to 10% for the remainder of the protocol.

### Tat-Beclin1 administration

Tat-Beclin1 was obtained from Novus Biologicals (Cat No. NBP2-49888). Tat-Beclin1 was reconstituted in sterile 1x phosphate-buffered saline (PBS). Mice in the TB and TB+Exe groups received intraperitoneal injections of 15 mg/kg of Tat-Beclin1 on Tuesdays and Thursdays, totaling 8 injections (Figure 1). Control and Exe groups were given PBS as a control for drug injection and to account for stress from the animal handling procedure. The chosen Tat-Beclin1 dose was supported by studies highlighting its efficacy as an autophagic inducer (26, 31, 32).

### Protein extraction, western blotting, and antibodies

Forty-eight hours after the last TB injection or the last exercise session, gastrocnemius/plantaris (G/P) and tibialis anterior (TA) muscles were harvested, weighed, flash-frozen in liquid nitrogen, and stored at – 80°C. Cryo-pulverized G/P and TA muscles were homogenized in a mix of ice-cold RIPA buffer (Thermo Fisher Scientific, Waltham, MA; Cat. No. 89901) and protease and phosphatase inhibitor cocktail (Thermo Fisher Scientific, Waltham, MA; Cat. No. 78441). Homogenates were centrifuged at 15,000 g for 15 min at 4 °C. We used a BCA Protein Assay Kit (Thermo Fisher Scientific, Waltham, MA; Cat. No. 23227) to measure total protein concentration. Ten or fifteen micrograms of total protein were loaded in an SDS-PAGE gel and transferred to a 0.22 μm or 0.45 μm PVDF membrane. Membranes were blocked with 5% skim milk in 1x PBS + 0.1% Tween-20 for 1 hour at RT, followed by overnight incubation with primary antibodies in a similar solution. Primary antibodies used in this study include microtubule-associated protein 1 light chain 3B (LC3B, 1:1000, No. ab192890), Beclin1 (1:2000, No. ab207612) from Abcam (Cambridge, UK), Sequestosome1/p62 (SQSTM1/p62, 1:1000, No. 23214) from Cell Signaling Technology, and Orosomucoid-1 (ORM1, 1:1000, No. MA5-24279) and Haptoglobin (HP, 1:1000, No. MA5-32919) from Invitrogen. Membranes were washed 3x for 10 minutes each in 1x PBS + 0.1% Tween-20, then incubated with secondary antibodies diluted in 1x PBS + 0.1% Tween-20. Coomassie Brilliant Blue G 250 was used as a loading control and normalization method.

### Immunofluorescence

G/P and TA muscles were harvested, placed onto a cork, and their distal and proximal portions fixed with metal pins. Muscles were then submerged in liquid nitrogen-cooled isopentane for ∼30 seconds and stored at – 80°C. The following day, the cryostat was set to – 28°C; the muscles were trimmed with a razor blade at the distal and proximal portions, near the metal pins, and embedded in a thin layer of Optimal Cutting Temperature (O.C.T.®) by submerging the samples in liquid nitrogen-cooled isopentane for 10 seconds. After all samples were embedded in O.C.T.®, the cryostat was set to –21°C, and the G/P and TA muscles were cut at 10 μm, collected using Epredia™ Shandon™ ColorFrost™ Plus Slides (ThermoFisher), and stored at – 80°C until ready for staining. Next, slides containing G/P and TA were air-dried for 20 min at room temperature, washed in PBS for 5 min, blocked in 10% goat serum, 0.05% Triton X-100, and PBS for 1 hour at RT. After washing sections with PBS for 5 min, 3x each, samples were incubated for 2 hours at RT with SC-71 (mouse IgG1, 1:400) to detect myosin heavy chain IIa, and BF-F3 (mouse IgM, 1:50) to detect myosin heavy chain IIb, purchased from the Developmental Studies Hybridoma Bank, and Laminin (Ab11575, rabbit IgG, 1:100) from Abcam. The SC-71 and BF-F3 antibodies developed by Schiaffino, S., were obtained from the DSHB, created by the NICHD of the NIH, and maintained at The University of Iowa, Department of Biology, Iowa City, IA 52242. Following primary antibody incubation, muscle sections were washed with PBS (5 min, 3x), and incubated with secondary antibody as follows: Alexa Fluor 488 (MIgG1, 1:200), Alexa Fluor 555 (MIgM, 1:200), and Alexa Fluor 350 (RbIgG (H+L), 1:200) for 1 hour at RT. A drop of ProLong™ Gold Antifade Mountant was applied to the sections, which were then covered with a coverslip and sealed with clear nail polish. Slides were immediately imaged or stored at 4°C. Multichannel images of the G/P and TA muscle sections were captured using a Leica DM4000 B LED microscope equipped with a K3M camera. High-resolution images of the entire muscle section were assembled using the Grid/Collection stitching plugin in Fiji in sequential-image mode. Next, multichannel fluorescent images were segmented using Cellpose Sam, and the resulting masks were imported into the LabeltoROIs plugin in Fiji, alongside the original images, to accurately quantify myofiber features. The data generated by this process were subsequently analyzed using a custom Python script for detailed processing.

### Proteomics

Cryopulverized G/P muscle tissue was lysed, solubilized, and digested on an S-Trap Micro (Protifi). The resulting peptides were de-salted using an Oasis HLB µElution plate (Waters), dried via vacuum centrifugation, and reconstituted in 0.1% TFA containing iRT peptides (Biognosys Schlieren, Switzerland). Peptides were analyzed using a QExactive HF mass spectrometer (Thermo Fisher Scientific, San Jose, CA) coupled with an Ultimate 3000 nano UPLC system and an EasySpray source, employing data-independent acquisition (DIA). The raw mass spectrometry data were searched using direct-DIA mode in Spectronaut. The MS2 intensity values for proteins generated by Spectronaut were used for bioinformatics analysis.

### Statistical analyses

Data is presented as means ± standard deviation (SD). Outliers were identified using the Grubbs’ method (α = 0.05). The Shapiro-Wilk normality test was applied to assess whether the data were normally distributed; any non-normally distributed samples were excluded from the analysis. Repeated measures Two-way ANOVA with Šídák’s multiple comparisons test was applied for pre- and post-assessments. Two-way ANOVA with Tukey’s multiple comparisons test was applied to detect differences among groups. For these tests, an α < 0.05 was considered significant. Proteomics intensities were filtered to remove proteins with excessive missingness and normalized using sample-wise median centering to reduce technical variation. Differential protein abundance between groups was evaluated using linear modeling with empirical Bayes variance moderation as implemented in the limma framework, a widely used method for high-dimensional omics data that improves variance estimation and statistical power. Multiple testing correction was performed using the Benjamini–Hochberg (BH) procedure, and significantly differentially expressed proteins were subsequently examined using volcano plots and hierarchical clustering heatmaps. To assess pathway-level enrichment patterns, we performed Gene Set Enrichment Analysis (GSEA) using curated KEGG pathway gene sets. The results of the GSEA were summarized using a bar plot of NES values for significantly enriched pathways and Circular pathway plots. Proteins with P < 0.05, |log2 fold change| ≥ 1, and FDR < 0.1 were prioritized for downstream Gene Ontology and KEGG analyses.

## Results

### Impact of Tat-Beclin1 and endurance training on body and muscle weight

We assessed the mouse’s body weight pre- and post-treatment. Treating mice with TB resulted in a 12% decrease in body weight (BW) from pre-to post-treatment (Figure 2A, P = 0.003), whereas no difference was observed in the control, Exe, and TB+Exe groups. While all groups demonstrated a drop in body weight (Figure 2B), the reduction in BW in the TB group resulted in a significant net loss compared to the control (P = 0.0092) and TB+Exe groups (P = 0.0226). In addition, we found no significant differences in the muscle masses of G/P, TA, extensor digitorum longus, and soleus (Figures 2C-F; P > 0.05).

**Figure 2.**
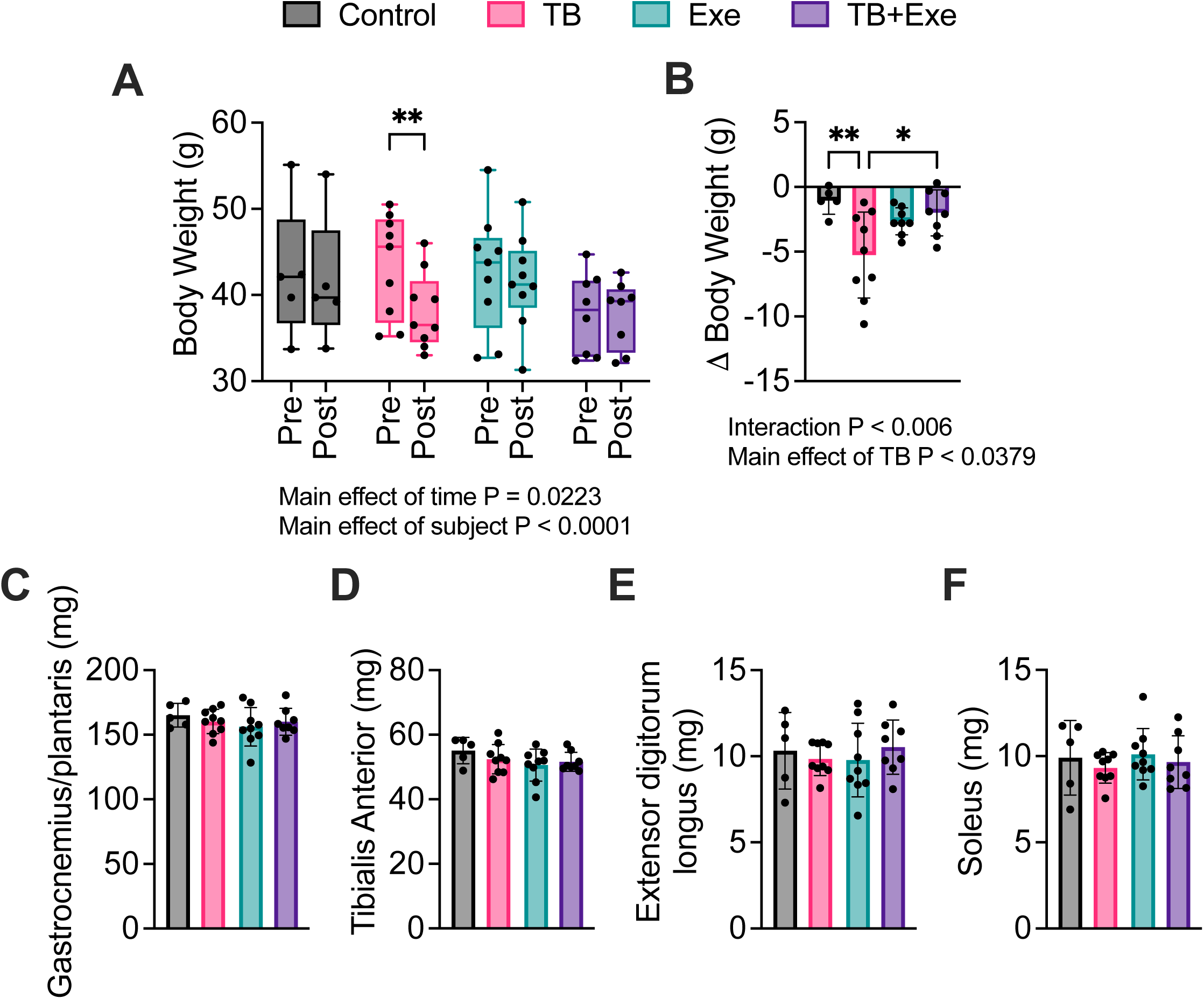
Body weight and hindlimb muscle masses following Tat-Beclin1 and endurance training. (A) Body weight (g) was measured before and after the intervention in Control, Tat-Beclin1 (TB), Exercise (Exe), and TB+Exe groups. (B) Change in body weight (Δ body weight, g). (C–F) Wet muscle masses for gastrocnemius/plantaris (C), tibialis anterior (D), extensor digitorum longus (E), and soleus (F). Data are presented as mean ± standard deviation with individual data points shown. For body weight measured pre- and post-intervention, statistical analysis was performed using two-way repeated-measures ANOVA followed by Šídák’s multiple comparisons test. Δ body weight and muscle masses were analyzed by two-way ANOVA followed by Tukey’s multiple comparisons test. Significant effects shown in the figure include a main effect of time for body weight (P = 0.0223), a main effect of subject (P < 0.0001), an interaction for Δ body weight (P < 0.006), and a main effect of TB (P = 0.0379). 23-month-old male C57BL6/J mice: Control, N = 5; TB, N = 9; Exe, N = 9; TB+Exe, N = 8.

### Tat-Beclin1 alone improves physical function, but Tat-Beclin1 and endurance training further enhance it

To determine whether TB, or its addition to a short-term endurance training regimen, affects physical function in old male mice, we measured grip strength, endurance capacity on a treadmill (time to exhaustion), and balance and coordination on a rotarod. Evaluating the effects of the one-month intervention on grip strength, our data revealed that TB and TB+Exe groups increased grip strength post-treatment compared with pre-treatment (Figure 3A; +20%, P = 0.0056; +41%, P < 0.0001, respectively), whereas no difference was observed in the control or Exe groups. The net gain was higher in TB and TB+Exe (Figure 3B, P = 0.0150, P = 0.0004, respectively) than in the control group, and in TB+Exe (P = 0.0198) than in the Exe group. Next, we conducted an incremental test to assess endurance capacity. Compared with pre-treatment, mice in the TB, Exe, and TB+Exe groups improved their endurance capacity (Figure 3C; TB +36%, P = 0.0025, Exe +53%, P < 0.0001, and TB+Exe +66%, P < 0.0001, respectively).

**Figure 3.**
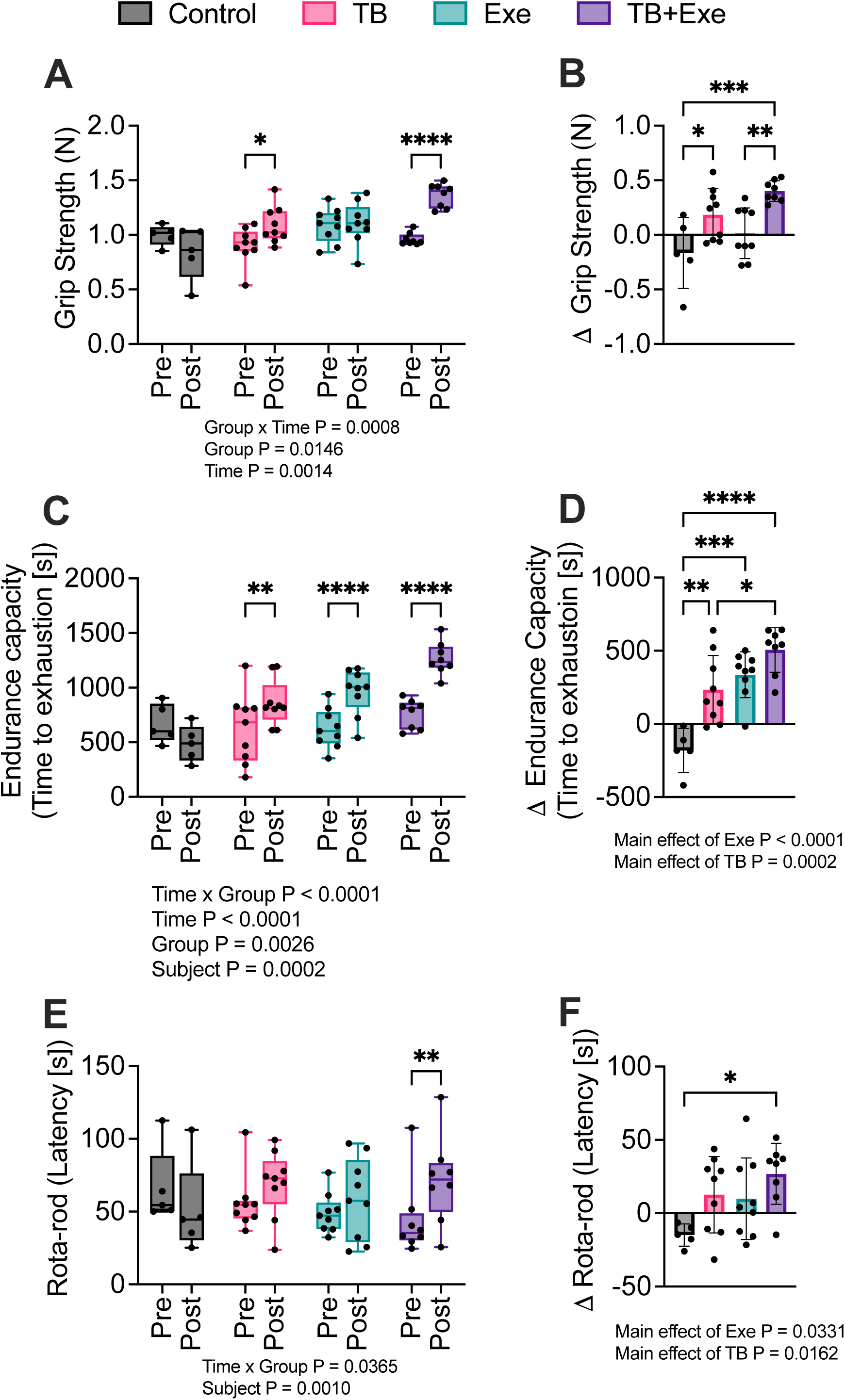
Tat-Beclin1 enhances muscle strength and endurance and induces additive effects when combined with endurance training in aged mice. (A) Four-paw grip strength (N) was measured before and after the intervention in Control, Tat-Beclin1 (TB), Exercise (Exe), and TB+Exe groups. (B) Change in grip strength (Δ grip strength, N). (C) Endurance capacity, reported as time to exhaustion (s), was measured before and after the intervention. (D) Change in endurance capacity (Δ endurance capacity, s). (E) Rotarod performance, reported as latency to fall (s), was measured before and after the intervention. (F) Change in rotarod performance (Δ rotarod latency, s). Data are presented as mean ± standard deviation with individual data points shown. Pre- and post-intervention measures were analyzed using two-way repeated-measures ANOVA followed by Šídák’s multiple comparisons test. Δ grip strength, Δ endurance capacity, and Δ rotarod latency were analyzed using two-way ANOVA followed by Tukey’s multiple comparisons test. Significant effects shown in the figure include group × time interactions for grip strength (P = 0.0008), endurance capacity (P < 0.0001), and rotarod latency (P = 0.0365). Main effects shown include group (grip strength, P = 0.0146; endurance capacity, P = 0.0026), time (grip strength, P = 0.0014; endurance capacity, P < 0.0001), subject (endurance capacity, P = 0.0002; rotarod, P = 0.0010), Exe (Δ endurance capacity, P < 0.0001; Δ rotarod, P = 0.0331), and TB (Δ endurance capacity, P = 0.0002; Δ rotarod, P = 0.0162). 23-month-old male C57BL6/J mice: Control, N = 5; TB, N = 9; Exe, N = 9; TB+Exe, N = 8.

Compared to the control group, the net gain was increased in all three treated groups (Figure 3D; TB, P = 0.0018; Exe, P = 0.0001; and TB+Exe, P < 0.0001). In addition, the combination approach increased the net gain compared to the TB group (P = 0.0223). A main effect for Exe (P < 0.0001) and TB (P = 0.0002) was also noted. Because aging increases the risk of falls and fractures, we sought to determine whether our short-term therapeutic approach improves mice’s balance and coordination by conducting a rotarod assay. Compared with the pre-treatment period, only the TB+Exe group showed a 60% increase in post-treatment latency at the fall assessment, suggesting that TB+Exe might benefit subjects at risk of falls (Figure 3E, P = 0.0032). Additionally, the combination treatment yielded a greater net gain than the control group (Figure 3F, P = 0.0209).

### Impact of Tat-Beclin1, exercise, and their combination on the gastrocnemius/plantaris and tibialis anterior myofiber profile

To determine whether the short-term TB+Exe alters myofiber size and type in the G/P and TA muscles, we examined muscle sections processed for myosin heavy chain (MyHC) types IIa and IIb, and laminin. There was no difference in the G/P myofiber cross-sectional area (CSA) (Figures 4A and B, P > 0.05), in the CSA of each myofiber type (Figures 4A and C, P > 0.05), and in the G/P myofiber distribution (Figures 4A and C, P > 0.05) among the groups. Similar to the G/P muscles, the TA muscle showed no statistical differences for myofiber CSA, CSA of each myofiber type, and myofiber distribution (Figures 4F-I, P > 0.05).

**Figure 4.**
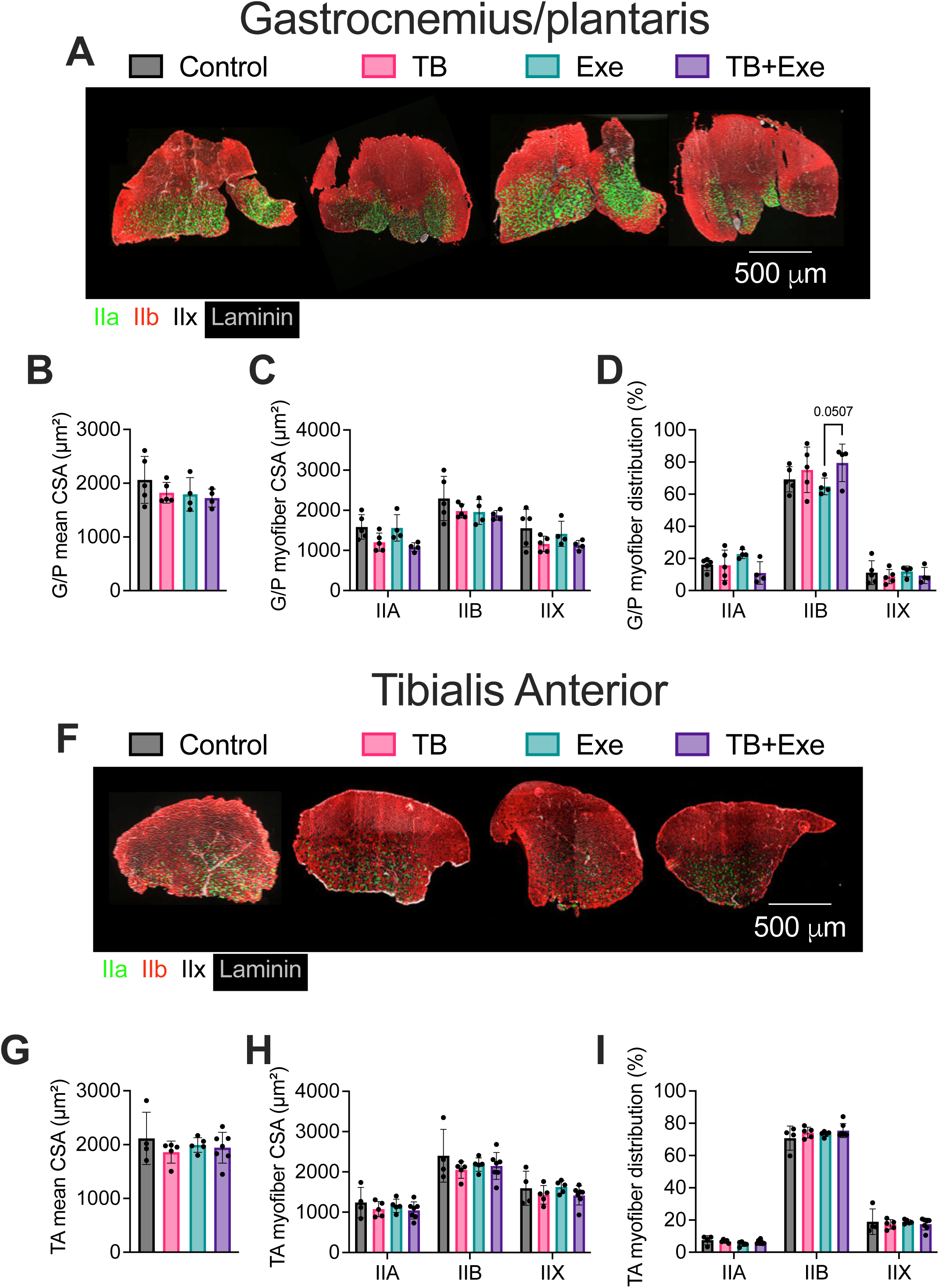
Gastrocnemius/plantaris and tibialis anterior fiber-type composition and cross-sectional area following Tat-Beclin1 and endurance training. (A) Representative gastrocnemius/plantaris cross-sections stained for myosin heavy chain fiber types, with type IIa fibers shown in green, type IIb fibers in red, type IIx fibers unstained/black, and laminin in gray outlining myofiber borders; scale bar, 500 µm. (B) Mean gastrocnemius/plantaris myofiber cross-sectional area (CSA, µm²). (C) Gastrocnemius/plantaris myofiber CSA by fiber type (IIa, IIb, IIx). (D) Gastrocnemius/plantaris myofiber-type distribution (%). (F) Representative tibialis anterior cross-sections stained as described above; scale bar, 500 µm. (G) Mean tibialis anterior myofiber CSA (µm²). (H) Tibialis anterior myofiber CSA by fiber type (IIa, IIb, IIx). (I) Tibialis anterior myofiber-type distribution (%). Data are presented as mean ± standard deviation with individual data points shown. Statistical analysis was conducted using two-way ANOVA. A trend is shown in gastrocnemius/plantaris fiber-type distribution (P = 0.0507). 23-month-old male C57BL6/J mice: Control, N = 4; TB, N = 5; Exe, N = 5; TB+Exe, N = 5.

### Protein expression of autophagy markers remains unchanged in the gastrocnemius/plantaris and tibialis anterior muscles after TB, Exe, or TB+Exe interventions

Given that our therapeutic approach combined Tat-Beclin1 and endurance training, two well-known autophagic agonists (23, 26), we assessed autophagy markers by western blotting in the G/P and TA muscles. Surprisingly, we found no obvious difference in the protein expression of the autophagic markers assessed, including Beclin1, LC3B-II/I ratio, and p62, in the G/P (Figures 5A-D) and TA (Figures 5E-H) muscles.

**Figure 5.**
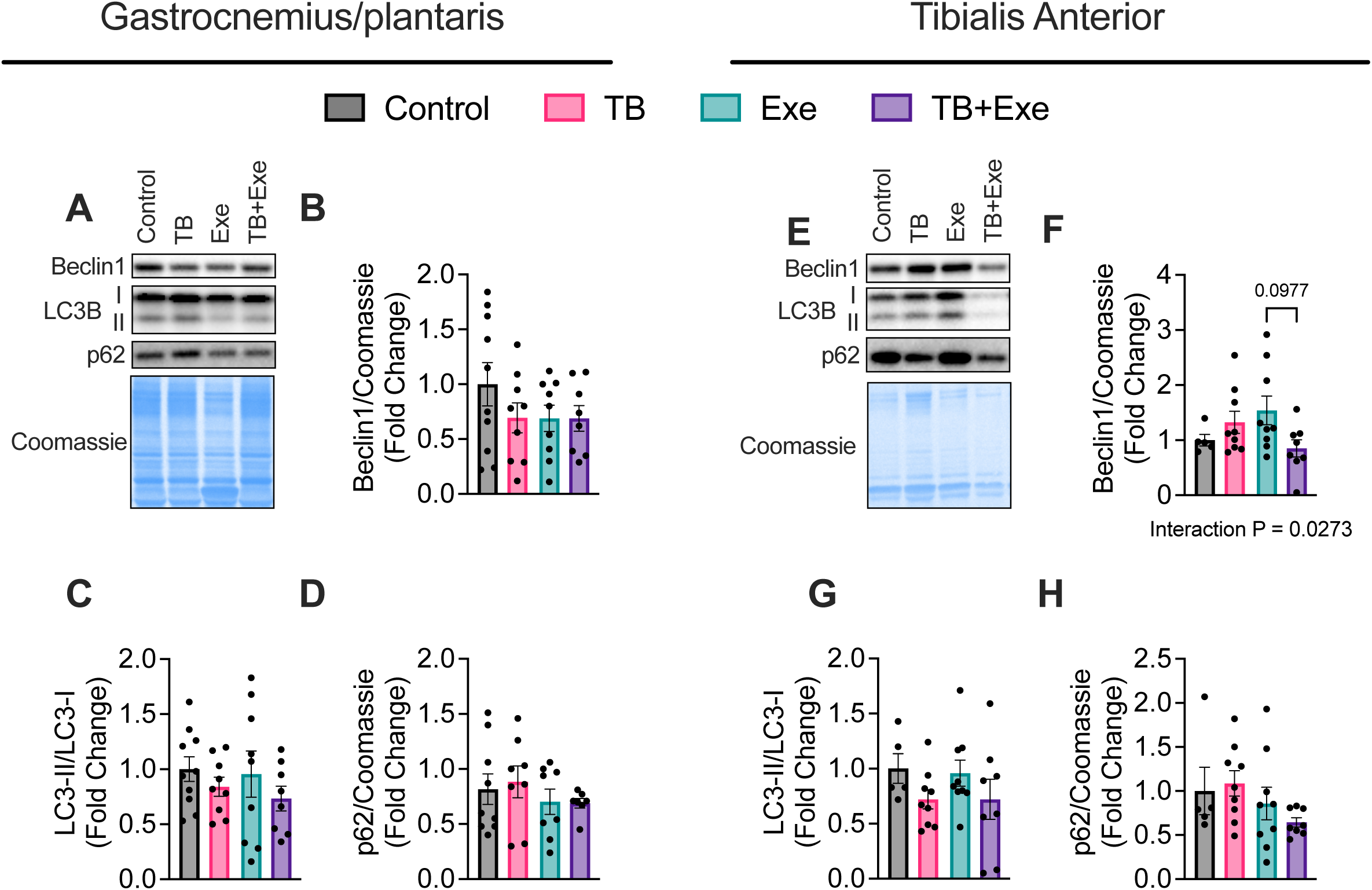
Autophagy-marker protein abundance in gastrocnemius/plantaris and tibialis anterior after Tat-Beclin1 and endurance training. Top panel, gastrocnemius/plantaris muscle: (A) Representative immunoblots for Beclin1, LC3B, and p62, with Coomassie brilliant blue staining shown as a loading control. Quantification is shown for (B) Beclin1/Coomassie, (C) LC3-II/LC3-I, and (D) p62/Coomassie. Right panel, tibialis anterior muscle: (E) Representative immunoblots for Beclin1, LC3B, and p62, with Coomassie brilliant blue staining shown as a loading control. Quantification is shown for (F) Beclin1/Coomassie, (G) LC3-II/LC3-I, and (H) p62/Coomassie. Data are presented as mean ± standard deviation with individual data points shown. Statistical analysis was conducted using two-way ANOVA. In tibialis anterior, Beclin1 showed a significant interaction (P = 0.0273) with a trend for pairwise comparison (P = 0.0977). 23-month-old male C57BL6/J mice: Control, N = 9; TB, N = 9; Exe, N = 9; TB+Exe, N = 8.

### Impact of combining Tat-Beclin1 and endurance training on the gastrocnemius/plantaris muscle

To investigate the mechanisms underlying functional adaptations induced by TB+Exe, we performed a proteomic analysis of G/P muscles. TB, Exe, and TB+Exe approaches showed distinct functional enrichment in Gene Ontology (GO) biological processes (GO: BP) and KEGG pathways compared with the control group (Suppl Figures S1A-F). The heatmap comparing the TB versus Exe groups showed a distinct protein pattern (Figure 6A). The GO: BP annotation showed upregulation of several muscle biology processes in the TB group versus the Exe group, including proteins involved in muscle contraction and adaptation. For instance, TB increased the levels of troponin proteins, including Tnnc1, Tnni2, and Scn4a, indicating changes in the regulation of muscle contraction (Figure 6B). The KEGG pathway analysis showed upregulation of proteins involved in energy pathways, including fatty acid and glutathione metabolism, as well as axon guidance (Figure 6C). When we compared TB+Exe with TB (Figures 6D-F), 1/3 of the proteins were upregulated in the TB+Exe group, as shown in the heatmap in Figure 6D. The annotated GO: BP revealed that the top 10 upregulated processes were related to mitochondrial bioenergetics, including cellular respiration, OXPHOS, ATP synthesis coupled to the electron transport chain, suggesting high aerobic energy production to support muscle function in the TB+Exe group. Interestingly, while OXPHOS and axon guidance were highlighted in the KEGG analysis as well, KEGG identified an increase in the complement pathway resulting from the TB+Exe strategy versus TB (Figure 6F).

**Figure 6.**
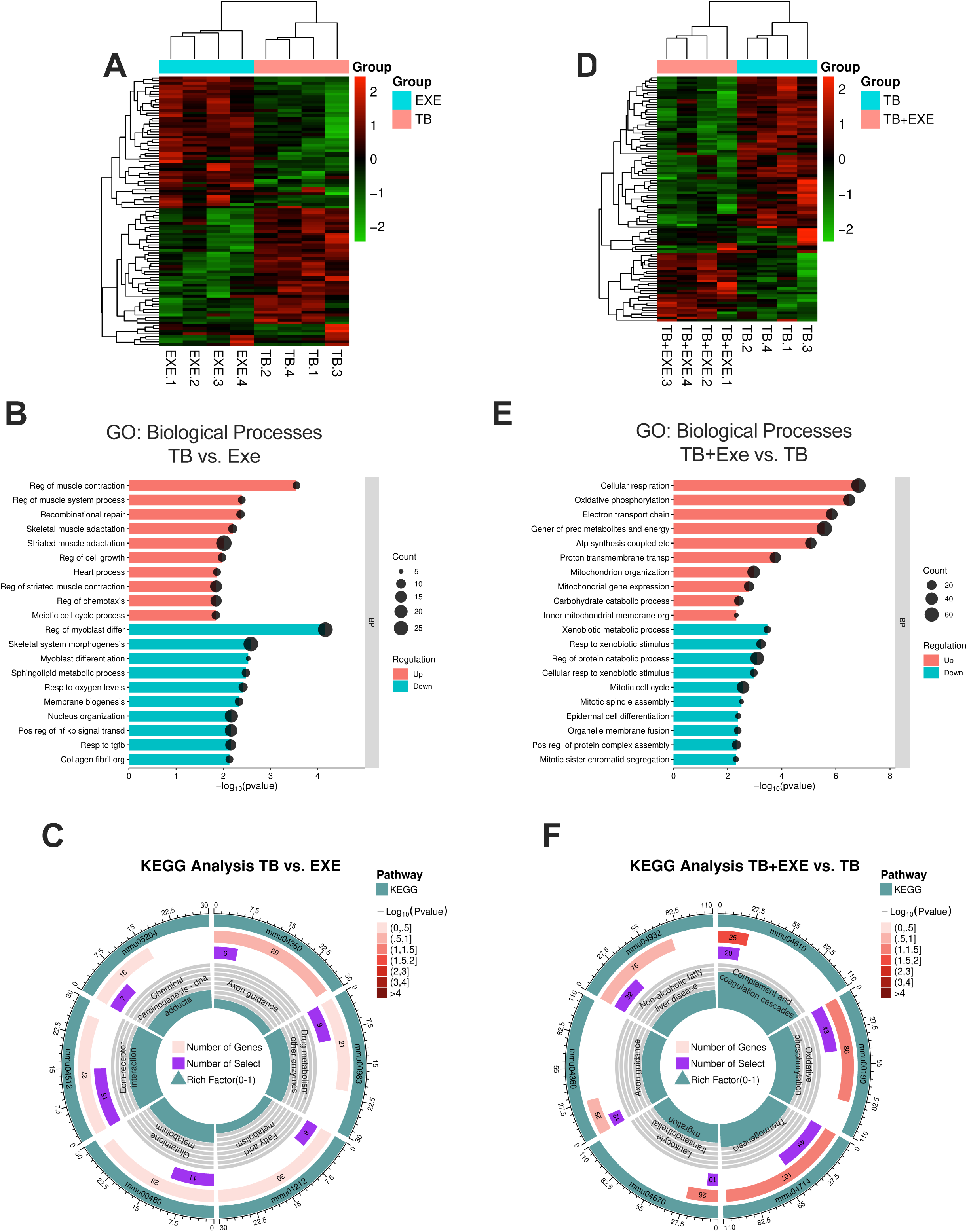
Proteomic signature of gastrocnemius/plantaris in TB vs. Exe and TB+Exe vs. TB. (A) Heatmap of the top 100 differentially abundant proteins comparing TB and Exe groups. (B) Gene Ontology (GO) Biological Process terms showing the top upregulated and downregulated processes, ranked by −log10(P value), in TB vs. Exe. (C) KEGG pathway analysis showing the top regulated pathways in TB vs. Exe. (D) Heatmap of the top 100 differentially abundant proteins comparing TB+Exe and TB groups. (E) GO Biological Process terms showing the top upregulated and downregulated processes, ranked by −log10(P value), in TB+Exe vs. TB. (F) KEGG pathway analysis showing the top regulated pathways in TB+Exe vs. TB. Analyses were corrected using the Benjamini-Hochberg false discovery rate (FDR < 0.1). 23-month-old male C57BL6/J mice used for proteomics: Control, N = 3; TB, N = 4; Exe, N = 4; TB+Exe, N = 4.

Moreover, the addition of Tat-Beclin1 to endurance training revealed that approximately 1/3 of the proteins were upregulated compared to the Exe group (Figure 7A). While the annotated functional enrichment in the GO: BP analysis for the TB+Exe group versus the Exe group indicated upregulation of proteins involved in OXPHOS and skeletal muscle contraction, it unexpectedly also showed upregulation of proteins involved in stress and acute inflammatory responses (Figure 7B). The KEGG pathway analysis revealed highly significant pathways involved in the organismal system (complement and coagulation), and the translational system (ribosome) (Figure 7C). Next, to identify the most statistically and biologically significant proteins in the GO: BP and KEGG analysis, we used a volcano plot (Figure 7D, P < 0.05 and Log_2_FC ≤ - 1 or ≥ 1). Of the 2154 proteins detected in the G/P muscle, only a small number met the statistical threshold. Peripherin (Prph), which is expressed in the neuromuscular junction, and mitochondrial ribosomal protein S7 (Mrps7), which is involved in protein synthesis, were both downregulated. Conversely, three proteins were upregulated in the TB+Exe group. We identified haptoglobin (Hp), which has been shown to be required to prevent muscle atrophy, inter-alpha-trypsin inhibitor heavy chain H4 (Itih4), which has a general function in tissue repair, and orosomucoid-1 (ORM-1), a protein shown to improve endurance capacity in young rats and mice (Figure 7D). This was further confirmed by WB analysis for Hp and ORM-1 in the G/P muscle of all four groups (Figure 7E). These proteins were chosen for their reported role in skeletal muscle atrophy and endurance (33, 34). Our data show that the Hp expression was increased in TB+Exe compared to Exe (Figure 7E and F; P = 0.0266) and a trend toward increased expression when compared to TB (Figure 7E and F; P = 0.0812). ORM-1 protein expression was increased in the TB+Exe group compared with the Exe group (Figure 7E and G; P = 0.0013) and showed a trend toward increased expression compared with the control (Figure 7E and G; P = 0.0803). Interestingly, TB treatment also increased ORM-1 expression compared to Exe (Figure 7E and G; P = 0.0171), indicating that the improvements in physical performance in TB groups may be partly driven by ORM-1.

**Figure 7.**
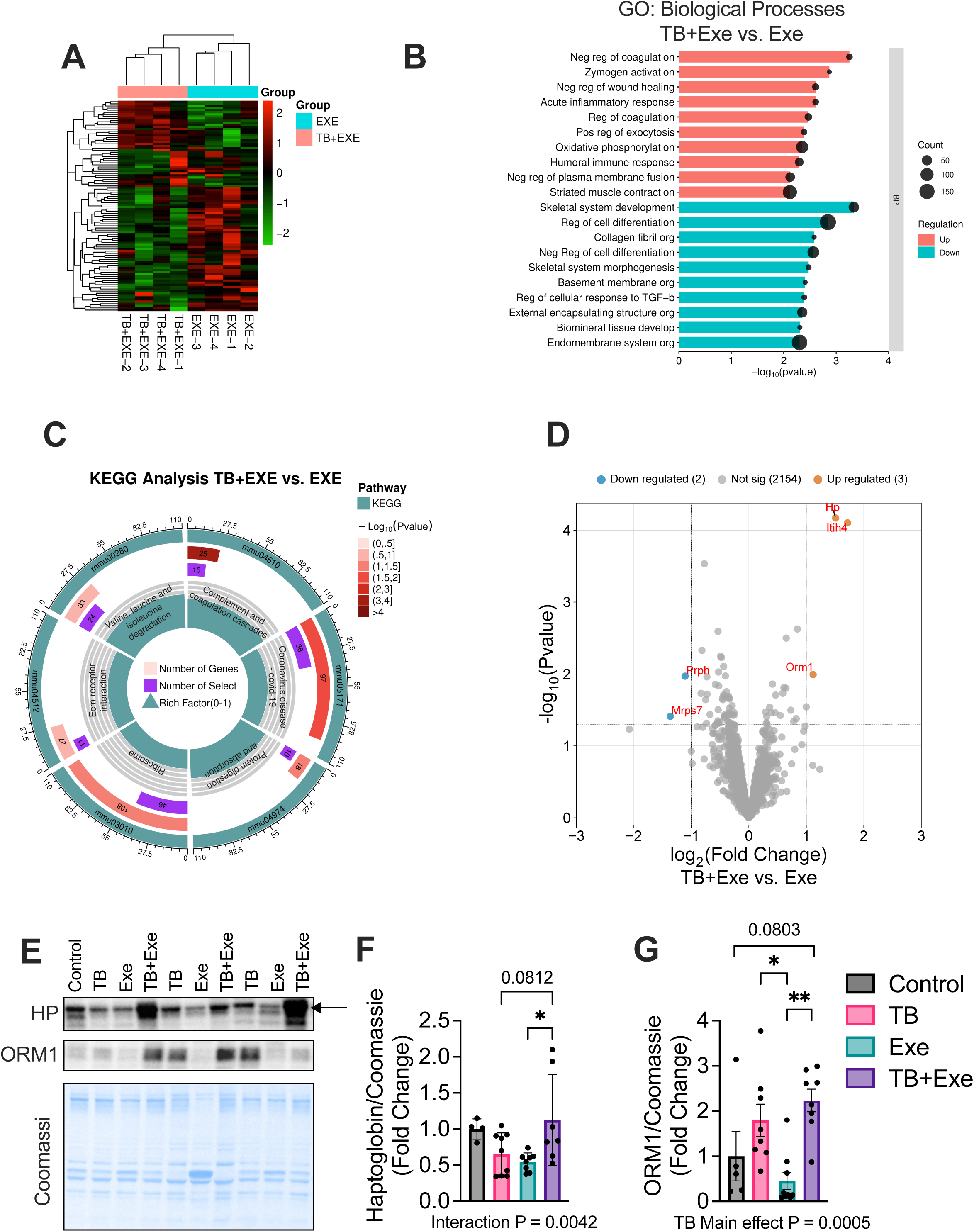
An acute-phase/inflammatory molecular signature distinguishes TB+Exe from Exe and validates induction of haptoglobin and ORM1. (A) Heatmap of the top 100 differentially abundant proteins comparing TB+Exe and Exe groups. (B) GO Biological Process terms showing the top upregulated and downregulated processes, ranked by −log10(P value), in TB+Exe vs. Exe. (C) KEGG pathway analysis showing the top regulated pathways in TB+Exe vs. Exe. (D) Volcano plot of TB+Exe vs. Exe showing log2 fold change versus −log10(P value), with significantly downregulated proteins shown in blue, significantly upregulated proteins shown in orange, and non-significant proteins shown in gray. Proteomics sample sizes: TB, N = 4; Exe, N = 4; TB+Exe, N = 4. (E) Representative immunoblots for haptoglobin (HP) and orosomucoid 1 (ORM1), with Coomassie brilliant blue staining as the loading control, across Control, TB, Exe, and TB+Exe groups. Quantification is shown for (F) HP/Coomassie and (G) ORM1/Coomassie. Data are presented as mean ± standard deviation with individual data points shown. Statistical analysis was conducted using two-way ANOVA followed by Tukey’s multiple comparisons test. For HP, an interaction was detected (P = 0.0042) with a trend for pairwise comparison (P = 0.0812). For ORM1, a main effect of TB was detected (P = 0.0005) with a trend for pairwise comparison (P = 0.0803). Immunoblot sample sizes: Control, N = 4-5; TB, N = 8; Exe, N = 9; TB+Exe, N = 7-8.

## Discussion

To the best of our knowledge, the current study is the first to investigate the impact of Tat-Beclin1 (TB), alone or in combination with endurance training, on the physical function of old male mice. The main findings of this study are that TB improves grip strength and endurance capacity relative to control animals, and that, strikingly, TB+Exe further increases gains in grip strength, endurance capacity, and coordination. In addition, proteomic profiling of the G/P muscle revealed that TB modulated proteins involved in muscle contraction, and that TB+Exe upregulated proteins associated with the acute inflammatory response. Our data suggest that this novel combinatorial strategy may enhance muscle protection against age-induced physical dysfunction, in part through an underexplored mechanism.

With advancing age, key physiological systems, including the cardiovascular, musculoskeletal, and nervous systems, begin to underperform, compromising the overall quality of life. In many preclinical studies, assessments of grip strength, time to exhaustion on a treadmill, and balance/coordination on a rotarod are used as affordable, rapid, and reliable methods to examine physical function (3, 28, 29, 35–37). In this study, 24-month-old male mice treated with TB increased four-paw grip strength and endurance capacity, whereas Exe increased only endurance capacity. Previous studies have shown promising outcomes with several agents in attenuating age-related declines in physical function (38–41). Among them, metformin, a widely used medication originally approved for treating type 2 diabetes, prevented muscle atrophy in older adults without diabetes after bed rest (14). Rapamycin, a potent mammalian Target of Rapamycin Complex 1 (mTORC1) inhibitor, improves grip strength, total running time, and neuromuscular junction function in aged mice (13). Additionally, an umbrella review of several pharmacological interventions concluded that only vitamin D (VitD) for older women with severely low VitD levels and testosterone for older men with combined muscle weakness and low levels of testosterone are clinically justifiable for attenuating age-related loss of muscle mass and function (i.e., sarcopenia) (15). Thus, the finding that TB improved grip strength and endurance capacity in aged mice indicates that a pharmacological approach targeting Beclin1 is a feasible strategy to attenuate age-related physical dysfunction.

While targeting aging-related physical dysfunction with a standalone pharmacological approach is promising, replicating these benefits when combining these agents with regular exercise is challenging (8). Metformin and rapamycin have been extensively studied in combination with regular exercise. When combined with progressive resistance exercise, metformin has been shown to antagonize the benefits of exercise in humans (16). In addition, metformin adversely affected aerobic exercise adaptation, primarily by attenuating muscle mitochondrial respiration in aged rodents and older adults (17, 18). Similarly, rapamycin opposes muscle hypertrophy adaptations to chronic overload and resistance exercise (19, 42). Intriguingly, the frequency of rapamycin administration seems to matter. A recent study (43) proposed two dosing schedules: 3x per week (frequent strategy) and 1x per week (intermittent strategy), both administered for 8 weeks in combination with progressive weighted wheel running (PoWer), a high-volume exercise modality shown to increase muscle mass and function (44). The study demonstrated that the intermittent strategy, combined with PoWer, produces promising outcomes in physical performance and muscle adaptations compared with a frequent-dosing strategy. Therefore, the present study and others suggest that careful planning of dosing strategies should be incorporated into experimental designs to maximize the drug-boosted increase in physical function when combined with regular exercise.

Endurance training and Tat-Beclin1 may independently activate autophagy (45, 46), yet our data show no change in the expression of autophagy markers (i.e., Beclin1, LC3B, and p62). A number of factors could have contributed to these outcomes in this study. For instance, the absence of assessment of LC3B-II turnover and p62 degradation by autophagic flux assays. Examination of autophagic flux can be achieved by adding a separate cohort of animals, reconducting the TB, Exe, and TB+Exe experiments, and ultimately treating them with lysosomal inhibitors (e.g., chloroquine) or agents that block autophagosome-lysosome fusion (e.g., colchicine) prior to euthanasia (47). Another possible explanation is the regulation of food intake prior to euthanasia, since rich nutrient conditions may blunt autophagic activity (48). In this study, we did not control the animals’ food intake, which may have masked alterations in the autophagy markers. Together, these factors suggest that the absence of changes in Beclin1, LC3B-II/I ratio, and p62 likely reflects limitations in detecting dynamic autophagic activity at the selected endpoint, rather than a true lack of autophagy induction by endurance training or Tat-Beclin1.

Next, we conducted a proteomic analysis to investigate the molecular changes associated with the improvements in physical function observed in the TB and TB+Exe groups. Compared with Exe, TB administration regulated several biological processes related to skeletal muscle contraction and organization, including upregulation of the troponin isoforms Tnnc1 and Tnni2, suggesting coordinated regulation of calcium binding and myosin-actin interactions. Also, the upregulation of proteins controlling cytoplasmic calcium during the contraction of slow-twitch and fast-twitch fibers, like Atp2a2 (SERCA2) and Atp2a1 (SERCA1), suggests highly orchestrated improvements in muscle function within MyHC type I and II. Studies have demonstrated that exercise increases troponin and SERCA expression (49, 50), suggesting that TB treatment improves physical function by activating pathways similar to those induced by traditional exercise training.

The combination of TB+Exe increased processes regulating mitochondrial energetics and the acute inflammatory response, as corroborated by the few proteins detected in the volcano plot and confirmed by the WB analyses. Hp and ORM1 are of particular interest in the context of skeletal muscle. Although they have several biological functions, ranging from drug binding and anti-sepsis to inhibiting superoxide production (ORM1) and from preventing excessive hemolysis to promoting antioxidant effects (Hp), both proteins have been previously linked to physical function and muscle atrophy. A 5-month-old Hp-knockout mouse model exhibited muscle atrophy, decreased muscle strength following an exercise challenge, and impaired physical performance (33), indicating the requirement for Hp to maintain physical function. ORM is emerging as an essential regulator of glucose and lipid metabolism (51).

Treating young rodents with ORM1 increased endurance capacity and muscle glycogen content (52). Additionally, in aged mice (28-month-old) subjected to 26 months of caloric restriction, known to activate autophagy, ORM1 was identified as a hub gene uniquely associated with caloric restriction (53). Mechanistically, it has been shown that ORM1 improves muscle endurance by activating AMPK, an energy sensor activating autophagy (34). However, the association between these proteins and autophagy remains elusive, at least in mammals, since ORM1 has been reported to be a selective autophagic receptor that interacts with ATG8 (LC3 ortholog) to activate autophagy in plants (54).

Collectively, our findings identify TB as a promising therapeutic agent to enhance physical function in aged male mice. They also show that an alternating-day regimen of TB+Exe yields additive functional benefits beyond either intervention alone. Proteomic profiling indicates that TB induces exercise-like adaptations in the muscle contractile apparatus. Meanwhile, TB+Exe underscores the involvement of mitochondrial bioenergetic pathways and an unexpected acute inflammatory response signature (e.g., Hp and ORM1), suggesting a potentially underexplored mechanism of muscle protection during age-related decline in physical function.

### Limitations of the present study and future directions

Our study provides new data that may pave the way for the inclusion of TB as a drug attenuating age-related physical dysfunction, especially when combined with endurance training. Nevertheless, we acknowledge a few limitations. We found no changes in autophagy markers following TB or TB+Exe treatment. Future studies should consider food removal for at least 6 hours prior to euthanasia. This would attenuate the confounding effects of recent food intake, since autophagy is finely regulated by nutrient status. Autophagy is a highly dynamic process, and adding a separate cohort treated with colchicine or chloroquine to assess autophagic flux should be considered in future studies. Moreover, given that age-related physical function deterioration is a slow, progressive process, future studies should determine whether long-term treatment with TB would elicit further improvements. Finally, prospective studies focusing on detected acute inflammatory proteins, including orosomucoid-1, are necessary to determine the benefits of TB+Exe and to assess whether it can serve as a standalone therapy for attenuating age-related decline in physical function.

## Acknowledgements

The authors thank Dhruti Brahmbhatt, Carina Davis, Maddy Murphy, and Saameh Siddique for their invaluable technical assistance in this study. This work was supported by the Camden Health Research Initiative (CHRI) and the NIH NIGMS award R16GM159126 and NIH NIA award R15AG095821 to KASS.

---

**Supplemental Figure S1. Gene Ontology Biological Process terms and KEGG pathway analysis for comparisons versus Control.**

(S1A, S1C, and S1E) GO Biological Process terms showing the top upregulated and downregulated processes, ranked by −log10(P value), in TB vs. Control, Exe vs. Control, and TB+Exe vs. Control, respectively. (S1B, S1D, and S1F) KEGG pathway analyses showing the top regulated pathways in TB vs. Control, Exe vs. Control, and TB+Exe vs. Control, respectively. Analyses were corrected using the Benjamini-Hochberg false discovery rate (FDR < 0.1). 23-month-old male C57BL6/J mice used for proteomics: Control, N = 3; TB, N = 4; Exe, N = 4; TB+Exe, N = 4.

