## Supplementary figures and images for "Short-Term Combined Tat-Beclin1 and Endurance Training Improves Age-Related Decline in Physical Function in Male Mice"

### Supplemental Figure 1

#### Supplemental Figure S1

S1A

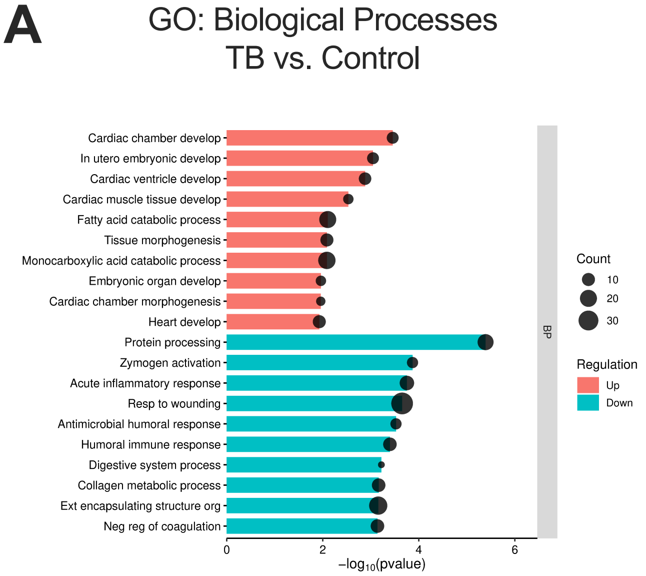


S1B

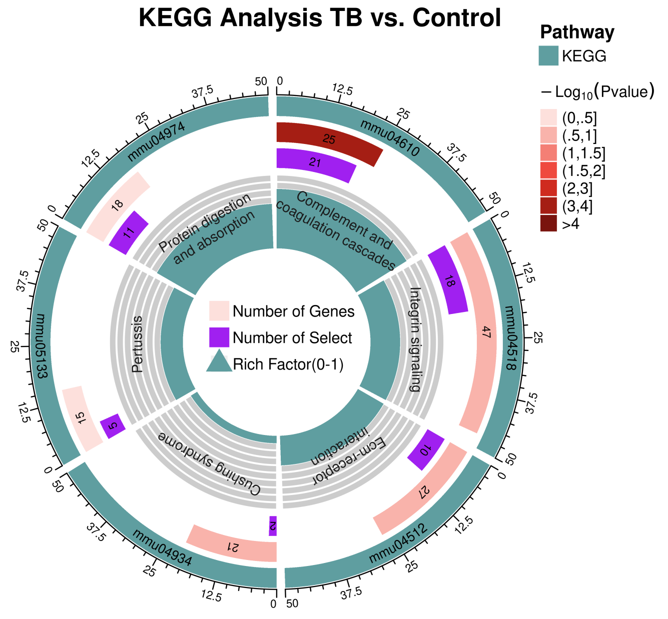


S1C

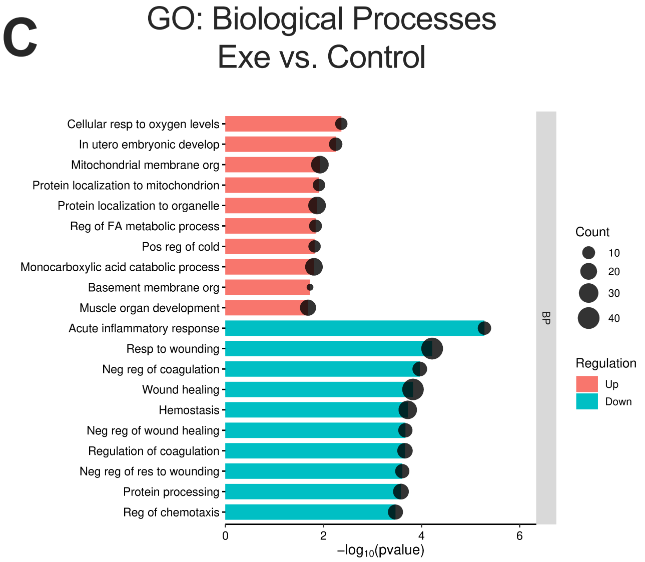


S1D

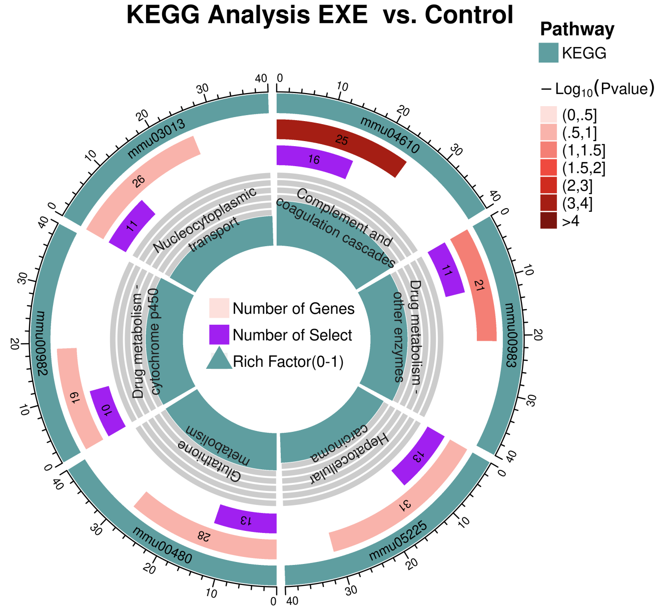


S1E

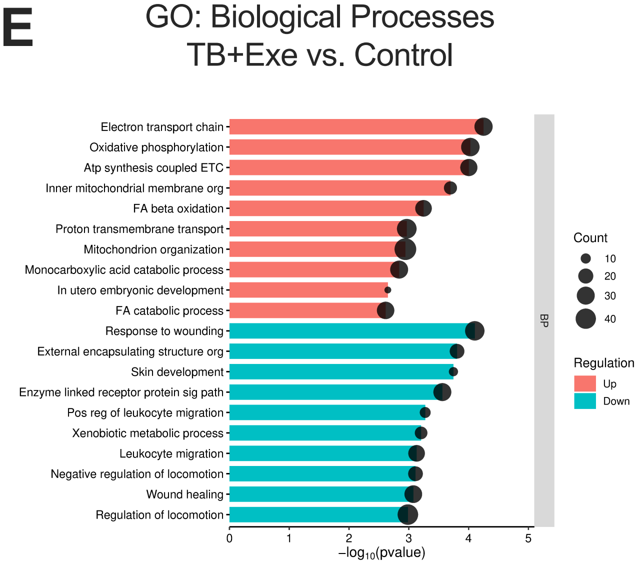


S1F

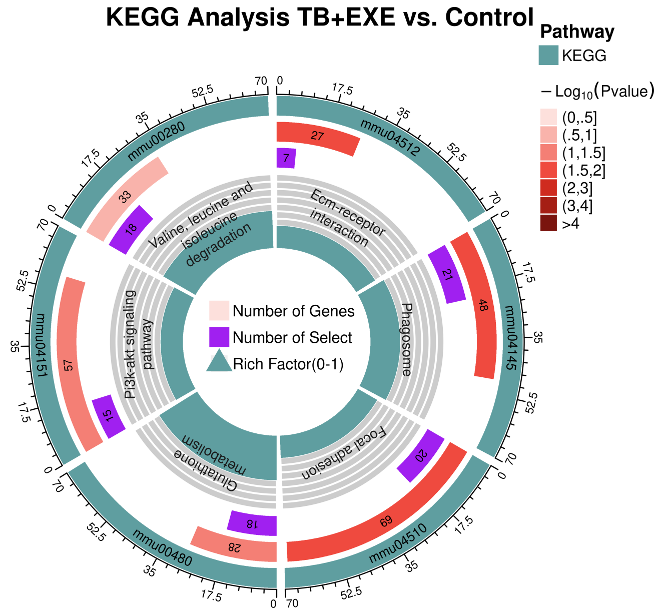
